# Integrated Transcriptomic and Network-Based Identification of Prognostic Hub Genes in Oral Squamous Cell Carcinoma

**DOI:** 10.64898/2026.04.02.716250

**Authors:** Sheilina Choudhary, Vandana Guleria

## Abstract

**Background:** The most prevalent kind of oral cancer is oral squamous cell carcinoma (OSCC), which has a poor prognosis because of delayed detection and a lack of molecular indicators.

**Methods:** Transcriptomic data from TCGA were analyzed to identify differentially expressed genes between OSCC and normal samples. Functional enrichment analysis was performed to determine biological pathways. A protein–protein interaction network was constructed using STRING and visualized in Cytoscape to identify hub genes.

**Results:** A total of 5732 differentially expressed genes were identified, including 2459 upregulated and 3273 downregulated genes. Network analysis revealed several highly connected hub genes such as CDK1, CCNB1, TOP2A, BUB1, and MMP9. Functional enrichment indicated significant involvement of cell cycle regulation and cancer-associated pathways.

**Conclusion:** This integrative analysis identified key regulatory hub genes that may be involved in OSCC progression. These genes may serve as promising biomarkers and therapeutic targets for future studies.

## 1. Introduction

Nearly 90% of all oral cancers globally are oral squamous cell carcinomas (OSCCs), the most common cancer of the mouth cavity. The prognosis for OSCC patients is still poor, with a five-year survival rate of roughly 50–60%, despite notable advancements in diagnostic and treatment methods(Bray et al., 2024). The main causes of the high death rate are aggressive tumor growth, delayed diagnosis, and a lack of trustworthy molecular biomarkers for focused treatment and early identification. The development of OSCC is a complex process that involves genetic, epigenetic, and environmental elements that work together to cause oral epithelial cells to undergo malignant transformation(Rivera, 2015; Warnakulasuriya, 2009).

The pathophysiology of OSCC has been linked to several risk factors, such as alcohol and tobacco use, poor oral hygiene, chronic inflammation, and infection-associated microbial dysbiosis(Warnakulasuriya, 2018). Oral potentially malignant disorders (OPMDs), which are precancerous lesions with the potential to develop into invasive carcinoma if left untreated, are quite prevalent, according to epidemiological research(Scully & Bagan, 2009). The need for better surveillance and early intervention strategies in high-risk populations is highlighted by a recent population-based study conducted in western Rajasthan, which revealed a high prevalence of oral potentially malignant disorders and identified several demographic and lifestyle determinants associated with disease occurrence. These results emphasize how crucial it is to comprehend the molecular pathways behind oral carcinogenesis to create efficient preventative and diagnostic strategies(Gautam et al., 2025).

Additionally, the oral microbiota may be crucial in regulating carcinogenic pathways, according to recent studies(Perera et al., 2016; Pleckaityte et al., 2012). In oral tissues, some pathogenic bacteria can affect host immune responses and encourage oxidative stress, persistent inflammation, and cellular damage. For example, it has been reported that the bacterial metabolite pyocyanin produced by Pseudomonas aeruginosa interacts with key host immune signaling components, such as NADPH oxidase and Toll-like receptor 4, potentially contributing to inflammatory signaling cascades linked to the development of oral cancer (Choudhary et al., 2025). These results imply that microbial metabolites may function as modulators of oncogenic signaling pathways, linking the pathophysiology of oral cancer to microbial dysbiosis.

Genetic changes and dysregulated gene expression largely influence OSCC progression in addition to environmental and microbiological variables(“Comprehensive Genomic Characterization of Head and Neck Squamous Cell Carcinomas,” 2015). Large-scale transcriptome profiling of cancer tissues has been enabled by advances in high-throughput sequencing, offering important insights into the genetic landscape of tumor formation. Differentially expressed genes (DEGs) that may have a role in the development, growth, invasion, and metastasis of tumors can be found using transcriptomic analysis(HUANG et al., 2015). However, the intricate relationships between cellular proteins and signaling networks involved in the development of cancer may not be adequately captured by examining gene expression alone(Hanahan & Weinberg, 2011).

Network-based bioinformatics techniques have become effective tools for identifying key regulatory genes in biological systems, thereby addressing this problem. Researchers can study functional relationships between proteins and find hub genes, highly linked nodes, by using protein–protein interaction (PPI) networks. These hub genes may be useful biomarkers or therapeutic targets since they frequently regulate vital cellular functions(Yu et al., 2007). Thus, a thorough foundation for comprehending the molecular pathways underlying cancer progression is provided by the integration of transcriptome data with PPI network analysis(Shannon et al., 2003; Szklarczyk et al., 2019).

Large-scale computational investigations of cancer transcriptomes have been made easier in recent years by the availability of vast datasets from publicly accessible genomic archives like The Cancer Genome Atlas (TCGA)(Weinstein et al., 2013). Researchers can methodically find dysregulated genes and pathways linked to tumor biology thanks to these databases. Key molecular fingerprints associated with the formation and progression of OSCC can be found by combining transcriptome profiling with functional enrichment analysis and protein interaction networks(Kanehisa et al., 2017; Tomczak et al., 2015).

Thus, the current study used an integrated transcriptomic and network-based bioinformatics method to discover differentially expressed genes and important prognostic hub genes associated with OSCC. Functional enrichment analysis was used to investigate related biological pathways after RNA-sequencing data revealed notable differences in gene expression between tumor and normal tissues. To find highly coupled hub genes that might be crucial to OSCC pathogenesis, a protein–protein interaction network was subsequently built. In addition to offering possible biomarkers for early detection and focused treatment approaches, the results of this study may advance knowledge of the molecular pathways driving oral cancer.

### 2. Materials and Methods

### 2.1 Data Acquisition and Preprocessing

The Cancer Genome Atlas (TCGA) database (https://www.genome.gov/Funded-Programs-Projects/Cancer-Genome-Atlas), specifically the Head and Neck Squamous Cell Carcinoma (HNSC) project, provided transcriptomic data for oral squamous cell carcinoma. RNA-sequencing gene expression patterns of tumor samples and matching normal tissue samples make up the dataset. The R statistical environment was used to retrieve and process the raw count data and related clinical metadata.

The data was pre-processed to eliminate low-quality or partial samples before downstream analysis. A matrix format was used to arrange the gene expression numbers, with rows denoting genes and columns denoting samples. To minimize technological variation and guarantee sample comparability, data processing and normalization were carried out.

### 2.2 Differential Gene Expression Analysis

To find genes that exhibited notable expression differences between OSCC tumor tissues and normal control samples, differential gene expression analysis was used. The R package DESeq2, which uses a negative binomial distribution to estimate variance and identify statistically significant differences in gene expression, was used in the investigation. To take into consideration variations in sequencing depth between samples, raw count data were first adjusted. Differential expression between the two groups was assessed using the Wald statistical test.

The Benjamini–Hochberg False Discovery Rate (FDR) correction method was used to modify p-values in order to account for multiple testing failures. Genes were deemed significantly differentially expressed if their absolute log2 fold change was larger than 1 and their adjusted p-value was less than 0.05. Graphical representations like scatter plots and volcano plots were used to display the findings of the differential expression study. The R tools ggplot2 and EnhancedVolcano were used for visualization. The volcano graphic makes it easy to identify upregulated and downregulated genes by highlighting those with notable fold changes and statistical significance.

### 2.3 Functional Enrichment Analysis

Functional enrichment analysis was used to determine the biological significance of the identified differentially expressed genes (DEGs). The genes were categorized using Gene Ontology (GO) analysis according to cellular components, molecular functions, and biological processes.

To find signaling pathways connected to the dysregulated genes, pathway enrichment analysis was also performed using the Kyoto Encyclopedia of Genes and Genomes (KEGG) database (https://www.genome.jp/kegg/pathway.html). The R package clusterProfiler was used to carry out these studies. A statistical criterion of adjusted p-value < 0.05 was used to identify significantly enriched GO keywords and pathways. The enrichment analysis’s findings shed light on important biological processes that contribute to OSCC formation.

### 2.4 Construction of Protein–Protein Interaction Network

A protein–protein interaction (PPI) network was constructed to examine functional interactions among the identified DEGs. The STRING database (https://string-db.org/), which offers information on known and anticipated protein interactions generated from computational predictions, experimental investigations, and public sources, received the list of important genes found in the differential expression study.

To filter interactions and guarantee network dependability, a confidence score threshold was used. Nodes in the resulting PPI network represented proteins, and interactions between them were represented by edges. This network offers a systems-level perspective on molecular interactions that could be involved in the pathophysiology of OSCC.

### 2.5 Network Visualization and Hub Gene Identification

For additional analysis and visualization, the PPI network created with STRING was imported into Cytoscape (https://cytoscape.org/). Key regulatory genes within the interaction network can be identified, and network structure can be thoroughly explored using Cytoscape.

In Cytoscape, hub genes were identified by analyzing topological metrics, such as node degree (the number of connections). Hub genes frequently play crucial roles in disease development and are regarded as important regulators within biological networks. For additional biological interpretation, the genes with the highest connection values were chosen as potential hub genes.

### 2.6 Statistical Analysis

The R programming environment was used for all statistical studies. When the adjusted p-value (FDR) was less than 0.05, differential expression results were deemed statistically significant. To ensure reproducibility and clarity of results, standard R packages were used to generate the plots and visualizations employed in the analysis.

## 3. Results

### 3.1 Identification of Differentially Expressed Genes

To identify genes associated with Oral Squamous Cell Carcinoma progression, differential gene expression analysis was performed using RNA-sequencing data obtained from the Cancer Genome Atlas database. After normalization and statistical filtering using the DESeq2 pipeline, a total of 5732 DEGs were identified between OSCC tumor samples and normal tissue samples.

Among these genes, 2459 genes were significantly upregulated, whereas 3273 genes were significantly downregulated in tumor samples compared with normal tissues. These results indicate widespread transcriptional alterations associated with OSCC development.

The distribution of significantly dysregulated genes was visualized using a volcano plot generated using Enhanced Volcano and ggplot2. In the volcano plot, as shown in **Figure 1**, genes with high fold change and strong statistical significance were clearly separated into upregulated and downregulated clusters, demonstrating distinct transcriptional patterns between cancerous and normal tissues.

**Figure 1:**
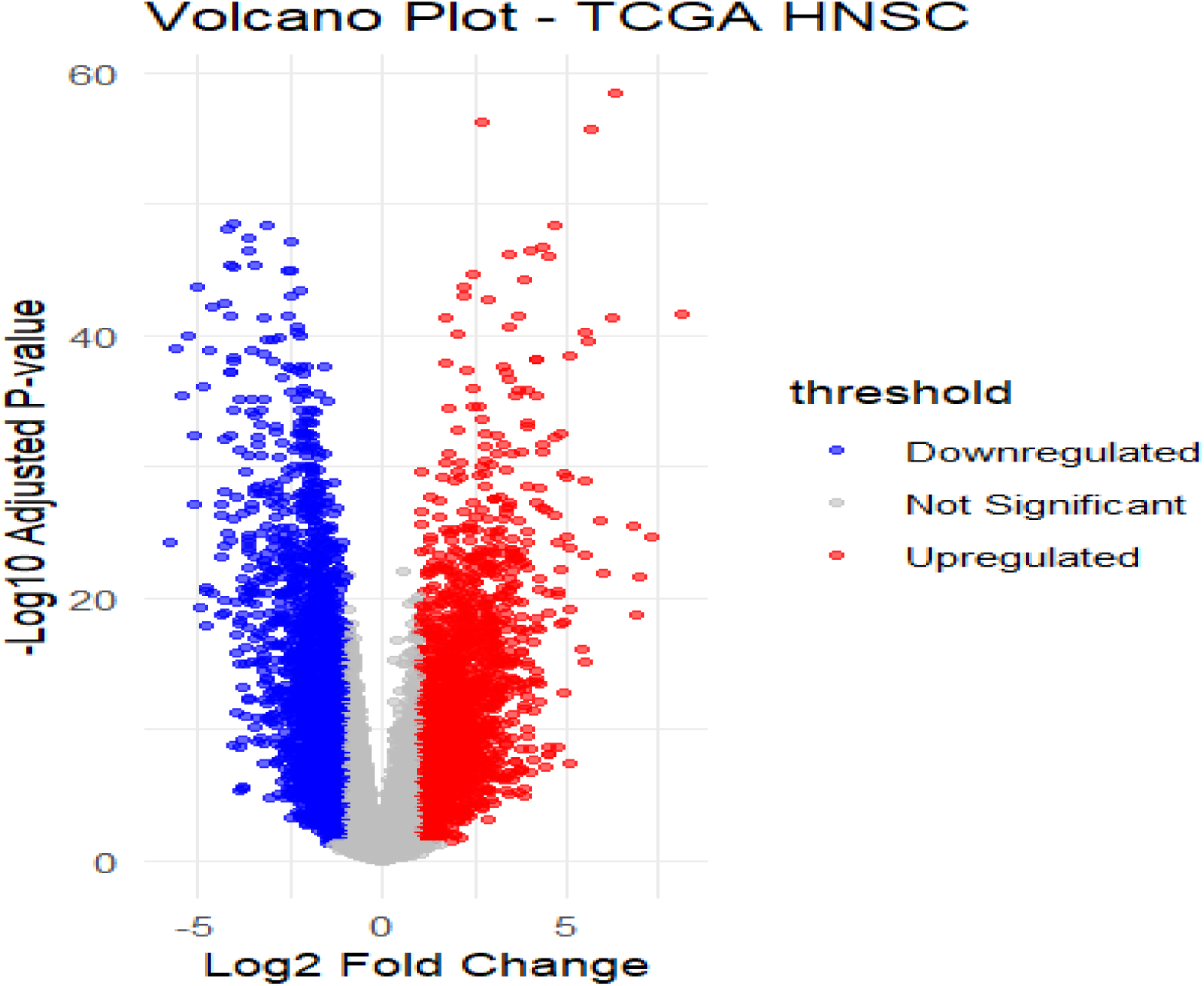
Volcano plot illustrating differentially expressed genes between OSCC tumor samples and normal tissues. The x-axis represents log2 fold change, while the y-axis represents –log10 adjusted p-value. Red dots indicate significantly upregulated genes, and blue dots represent significantly downregulated genes based on the defined threshold criteria.

### 3.2 Analysis of Functional Enrichment in Differentially Expressed Genes

Functional enrichment analysis was used to investigate the biological significance of the discovered DEGs. The clusterProfiler package in R was used to conduct pathway enrichment and Gene Ontology (GO) analyses.

Thousands of genes were evaluated concurrently in this Oral Squamous Cell Carcinoma investigation to find genes that were differently expressed between tumor and normal samples. Because transcriptome analysis involves several statistical tests, there is a substantial risk of receiving false positive results. As a result, the False Discovery Rate (FDR) was used to alter the p-values and increase the dependability of the identified genes, as explained in **Figure 2**.

**Figure 2:**
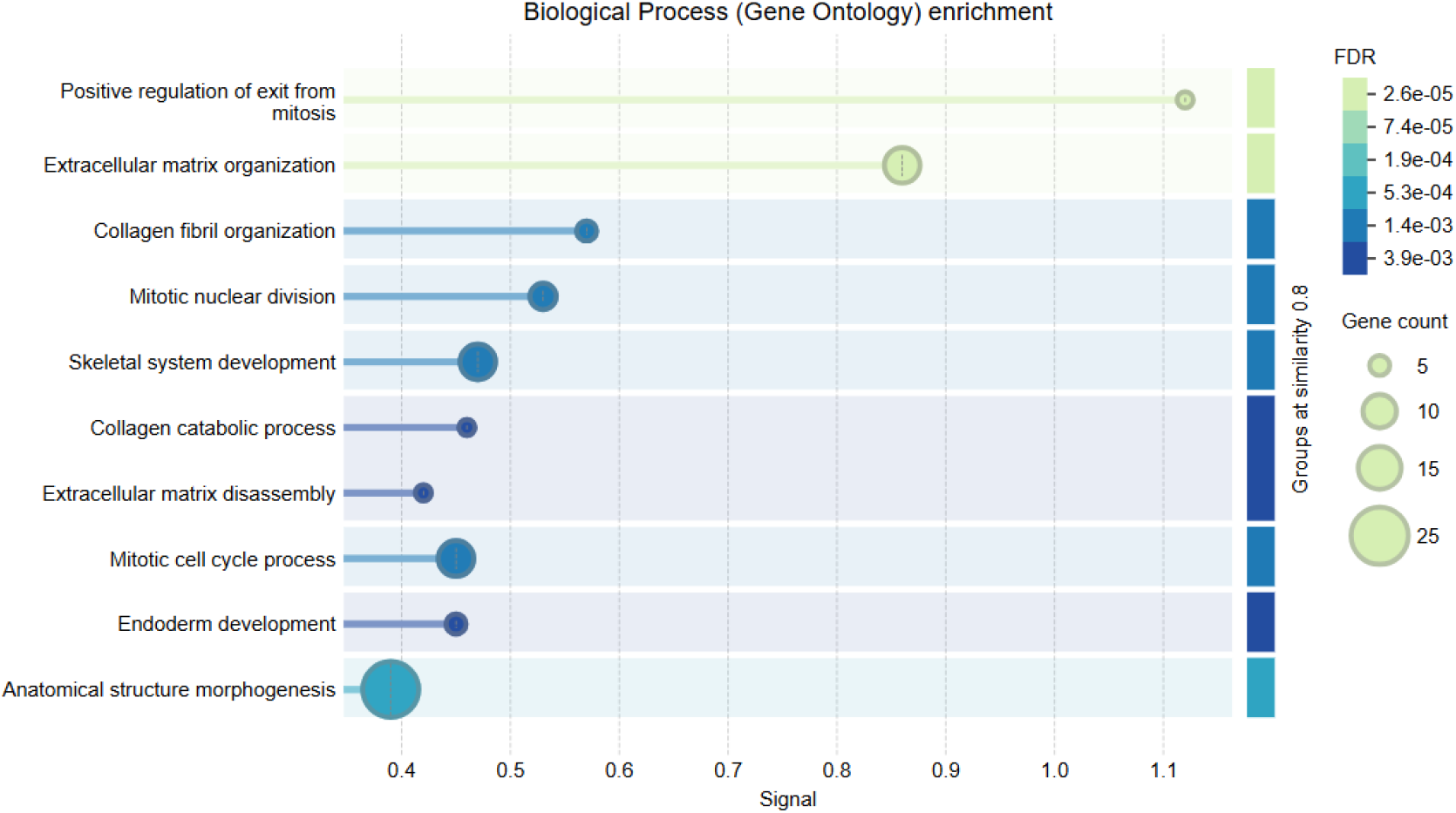
Gene Ontology (GO) Biological Process enrichment analysis of differentially expressed genes associated with Oral Squamous Cell Carcinoma. The bubble plot highlights significantly enriched biological processes, including extracellular matrix organization, collagen fibril organization, mitotic nuclear division, and morphogenesis of anatomical structures. Bubble size represents the number of genes involved in each process, while color intensity indicates the false discovery rate (FDR), reflecting the statistical significance of enrichment.

Using an FDR criterion (e.g., FDR < 0.05), only statistically significant and physiologically important genes were identified as differentially expressed. This guaranteed that the genes used in subsequent analyses, such as functional enrichment and protein-protein interaction network creation, were reliable and related to the molecular pathways that drive OSCC progression.

The analysis showed that biological processes associated with cell cycle regulation, mitotic cell division, extracellular matrix architecture, and cell proliferation were considerably enriched in the dysregulated genes. The development of tumors and the advancement of cancer are intimately linked to these basic processes.

Furthermore, pathway enrichment analysis identified several cancer-associated pathways, suggesting that the dysregulated genes are essential to the molecular processes underlying OSCC formation and tumor progression.

### 3.3 Construction of the Protein–Protein Interaction Network

Using the STRING database, a protein–protein interaction (PPI) network was built to examine the relationships between the discovered DEGs. To find known and anticipated interactions between their encoded proteins, the important genes were uploaded to the STRING platform. Numerous nodes, which represented proteins, and edges, which represented their functional relationships, made up the final PPI network. This interaction network, as shown in **Figure 3** enabled a systems-level understanding of the molecular connections among dysregulated genes implicated in OSCC. After that, Cytoscape was used to import the network for additional research and visualization. Cytoscape enabled the identification of highly connected nodes and a thorough investigation of the interaction network.

**Figure 3:**
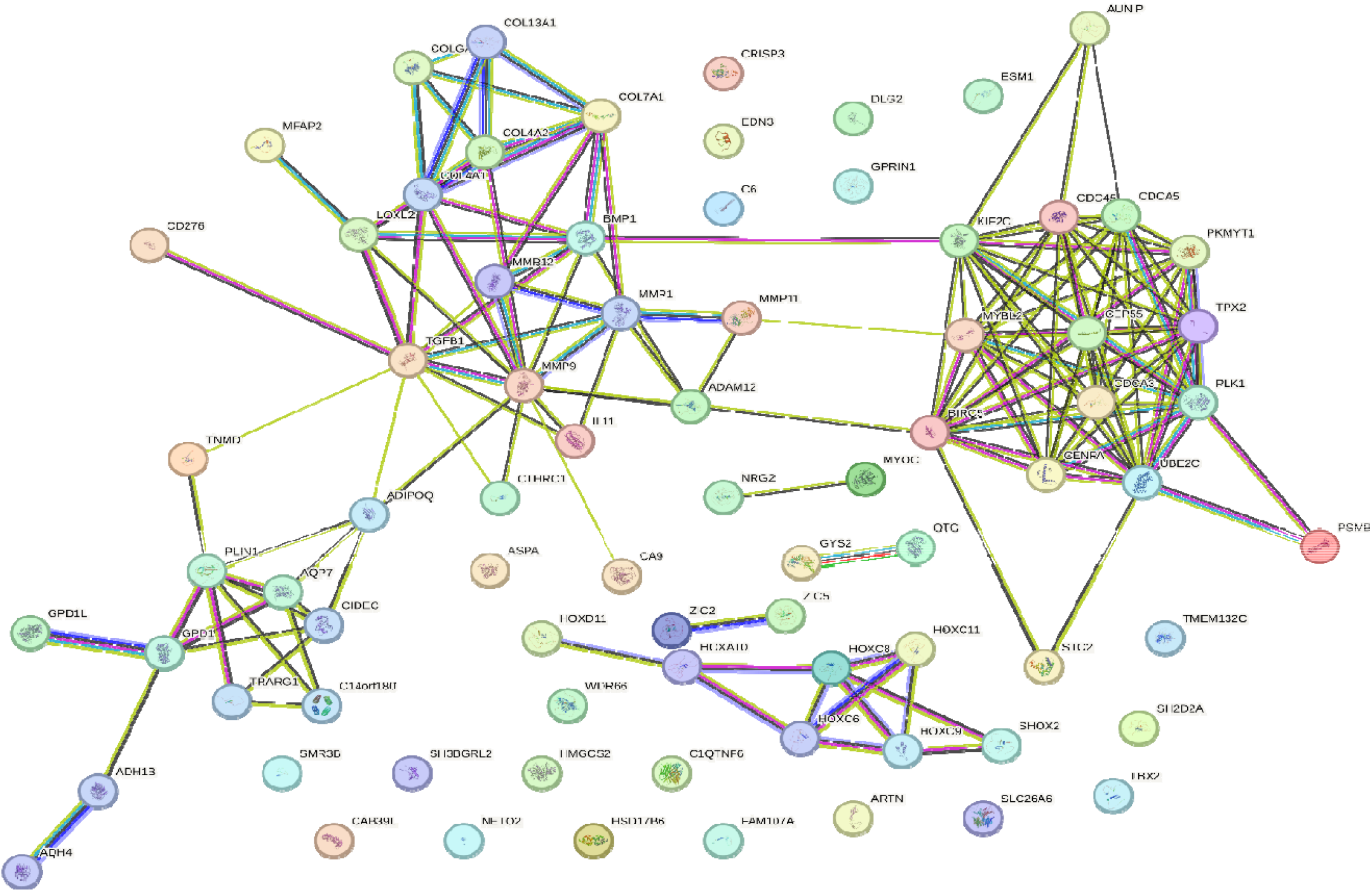
Protein–protein interaction (PPI) network constructed for the differentially expressed genes associated with Oral Squamous Cell Carcinoma using the STRING database and visualized in Cytoscape. Nodes represent proteins encoded by the identified genes, while edges indicate functional interactions between them, revealing key interaction clusters involved in OSCC progression.

### 3.4 Identification of Hub Genes

Additional Cytoscape network topology research identified several hub genes, or densely connected nodes, that may be important for the development of OSCC. Based on how connected they were inside the PPI network, hub genes were found. CDK1, CCNB1, TOP2A, BUB1, and MMP9 were shown to be important hub genes with strong connections to other proteins in the network, as shown in **Figure 4**. Important physiological functions such as DNA replication, cell cycle regulation, mitotic progression, and extracellular matrix remodelling are known to include these genes. These genes may function as key regulators in OSCC-associated biological pathways and may be useful biomarkers or therapeutic targets due to their significant interconnectedness.

**Figure 4:**
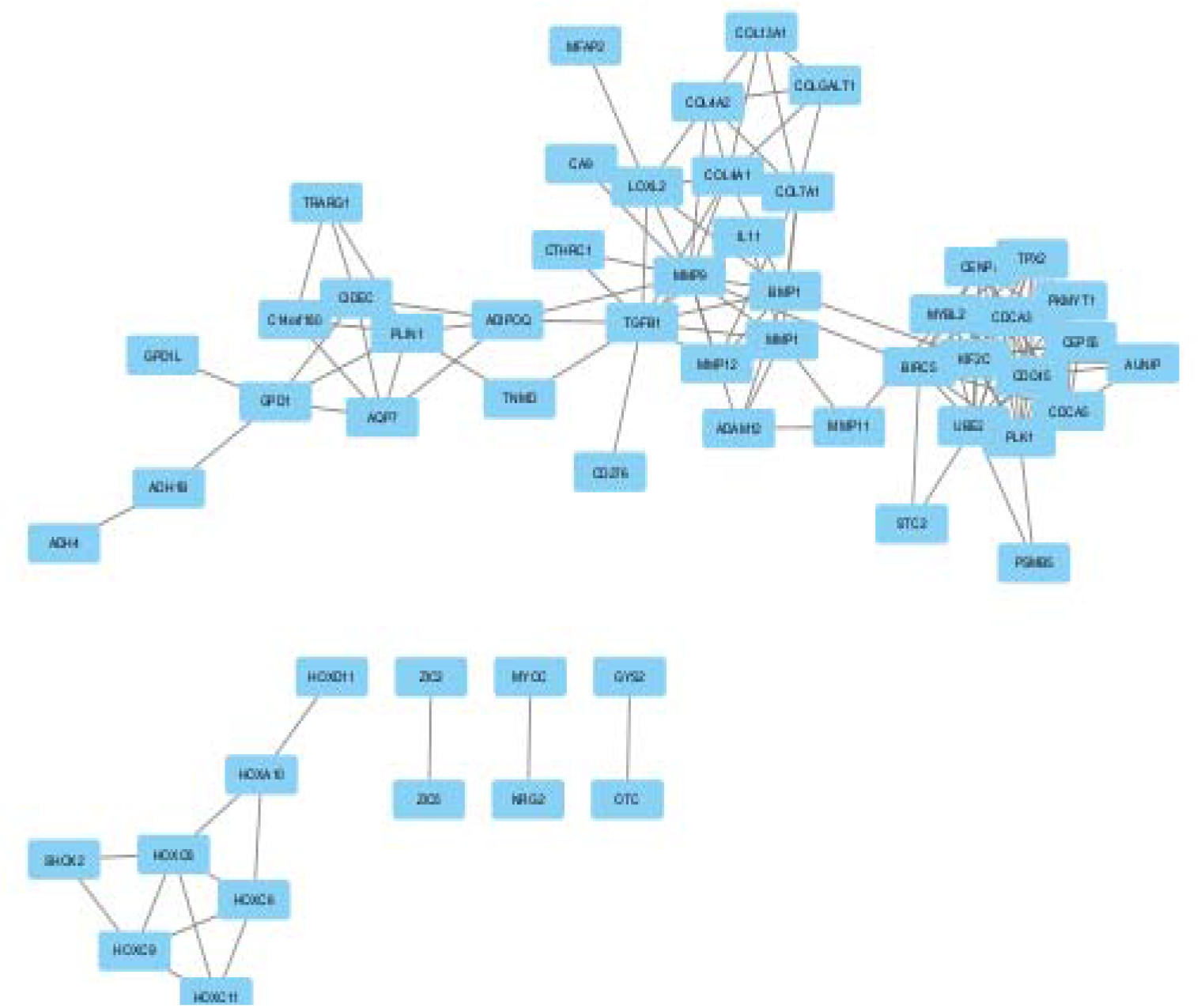
Hub gene interaction network identified from the protein–protein interaction (PPI) analysis of differentially expressed genes in Oral Squamous Cell Carcinoma. The network was constructed using the STRING database and visualized in Cytoscape. Hub genes were identified based on their high connectivity within the network, indicating their potential key roles in regulating molecular pathways involved in OSCC progression.

## 4. Discussion

The current work identified important genes and molecular interactions linked to oral squamous cell carcinoma using a thorough transcriptome analysis. When tumor tissues were compared to normal controls, differential gene expression analysis identified 5732 significantly dysregulated genes, including 2459 upregulated and 3273 downregulated genes. These results demonstrate widespread transcriptional changes that support the growth and evolution of OSCC.

The identified differentially expressed genes were found to be strongly linked to biological processes such as cell cycle regulation, mitotic cell division, extracellular matrix organization, and cellular proliferation, according to functional enrichment analysis. Since disruption of the cell cycle and aberrant cellular proliferation are hallmarks of the development of cancer, it is widely known that these biological processes play crucial roles in carcinogenesis.

A protein–protein interaction network was created using the STRING platform and then displayed in Cytoscape to investigate the functional interactions between these genes in more detail. A network topology study revealed some hub genes, highly linked nodes that may be crucial to the development of OSCC.

CDK1, CCNB1, TOP2A, BUB1, and MMP9 were among the hub genes shown to have substantial connectivity inside the interaction network. These genes are mostly involved in activities that are often dysregulated in cancer, such as DNA replication, cell cycle regulation, mitotic progression, and extracellular matrix remodeling. For example, abnormal expression of CDK1 and CCNB1, two important regulators of the G2/M phase of the cell cycle, is known to encourage unchecked cell division. Similar to how BUB1 is involved in the regulation of mitotic checkpoints, TOP2A is crucial for DNA replication and chromosomal segregation. Additionally, by breaking down extracellular matrix components and promoting cancer cell motility, MMP9 aids in tumor invasion and metastasis.

These hub genes may be useful biomarkers for early OSCC diagnosis and treatment targets, according to their discovery. Furthermore, the combination of network-based techniques with transcriptome analysis offers important insights into the intricate molecular pathways driving OSCC pathogenesis. Nevertheless, there are some drawbacks to this study. The results are mostly based on computational studies of transcriptome datasets that are accessible to the public, and experimental validation is necessary to verify the biological functions of the hub genes that have been identified. Further research into the functional importance of these genes and their possible therapeutic uses will require both in vitro and in vivo trials.

Overall, this study offers a comprehensive bioinformatics platform for identifying important genes and biochemical pathways involved in OSCC. The discovery of hub genes and related biological mechanisms may enhance our comprehension of OSCC progression and aid in the creation of innovative diagnostic and treatment approaches.

## 5. Conclusion

In order to uncover important genes linked to the advancement of oral squamous cell carcinoma, this study used an integrative transcriptomic and network-based method. 5732 substantially dysregulated genes were found by differential gene expression analysis, suggesting major molecular changes between tumor and normal tissues. Important biological processes that are intimately linked to the development of cancer, such as cell cycle regulation, mitotic division, and extracellular matrix architecture, were identified by functional enrichment analysis. CDK1, CCNB1, TOP2A, BUB1, and MMP9 were further identified by protein–protein interaction network analysis as putative hub genes that might be crucial to the pathophysiology of OSCC. These results offer insightful information on the molecular mechanisms driving OSCC and point to possible treatment targets and biomarkers for further study and clinical use.

## References

1. Bray, F., Laversanne, M., Sung, H., Ferlay, J., Siegel, R. L., Soerjomataram, I., & Jemal, A. (2024). Global cancer statistics 2022: GLOBOCAN estimates of incidence and mortality worldwide for 36 cancers in 185 countries. CA: A Cancer Journal for Clinicians, 74(3), 229–263. 10.3322/caac.21834

2. Choudhary, S., Kumar, P., Mohanty, S. S., Anand, P. K., & Gautam, J. K. (2025). Molecular docking of Pyocyanin from Pseudomonas aeruginosa with NADPH oxidase and Toll-like receptor 4L: Exploring the Link Between Oral Microbiome and Oral Cancer Pathogenesis. 10.1101/2025.08.24.671957

3. Comprehensive genomic characterization of head and neck squamous cell carcinomas. (2015). Nature, 517(7536), 576–582. 10.1038/nature14129

4. Gautam, J. K., Mohanty, S. S., & Babu, B. V. (2025). Prevalence and determinants of oral potentially malignant disorders in western rajasthan. Scientific Reports, 15(1), 15611. 10.1038/s41598-025-98096-8

5. Hanahan, D., & Weinberg, R. A. (2011). Hallmarks of Cancer: The Next Generation. Cell, 144(5), 646–674. 10.1016/j.cell.2011.02.013

6. Huang, G., Li, L., & Zhou, W. (2015). USP14 activation promotes tumor progression in hepatocellular carcinoma. Oncology Reports, 34(6), 2917–2924. 10.3892/or.2015.4296

7. Kanehisa, M., Furumichi, M., Tanabe, M., Sato, Y., & Morishima, K. (2017). KEGG: new perspectives on genomes, pathways, diseases and drugs. Nucleic Acids Research, 45(D1), D353–D361. 10.1093/nar/gkw1092

8. Perera, M., Al-hebshi, N. N., Speicher, D. J., Perera, I., & Johnson, N. W. (2016). Emerging role of bacteria in oral carcinogenesis: a review with special reference to periopathogenic bacteria. Journal of Oral Microbiology, 8(1), 32762. 10.3402/jom.v8.32762

9. Pleckaityte, M., Janulaitiene, M., Lasickiene, R., & Zvirbliene, A. (2012). Genetic and biochemical diversity of Gardnerella vaginalis strains isolated from women with bacterial vaginosis. FEMS Immunology & Medical Microbiology, 65(1), 69–77. 10.1111/j.1574-695X.2012.00940.x

10. Rivera, C. (2015). Essentials of oral cancer. International Journal of Clinical and Experimental Pathology, 8(9), 11884–11894.

11. Scully, C., & Bagan, J. (2009). Oral squamous cell carcinoma overview. Oral Oncology, 45(4–5), 301–308. 10.1016/j.oraloncology.2009.01.004

12. Shannon, P., Markiel, A., Ozier, O., Baliga, N. S., Wang, J. T., Ramage, D., Amin, N., Schwikowski, B., & Ideker, T. (2003). Cytoscape: A Software Environment for Integrated Models of Biomolecular Interaction Networks. Genome Research, 13(11), 2498–2504. 10.1101/gr.1239303

13. Szklarczyk, D., Gable, A. L., Lyon, D., Junge, A., Wyder, S., Huerta-Cepas, J., Simonovic, M., Doncheva, N. T., Morris, J. H., Bork, P., Jensen, L. J., & Mering, C.von. (2019). STRING v11: protein–protein association networks with increased coverage, supporting functional discovery in genome-wide experimental datasets. Nucleic Acids Research, 47(D1), D607–D613. 10.1093/nar/gky1131

14. Tomczak, K., Czerwińska, P., & Wiznerowicz, M. (2015). Review The Cancer Genome Atlas (TCGA): an immeasurable source of knowledge. Współczesna Onkologia, 1A, 68–77. 10.5114/wo.2014.47136

15. Warnakulasuriya, S. (2009). Global epidemiology of oral and oropharyngeal cancer. Oral Oncology, 45(4–5), 309–316. 10.1016/j.oraloncology.2008.06.002

16. Warnakulasuriya, S. (2018). Clinical features and presentation of oral potentially malignant disorders. Oral Surgery, Oral Medicine, Oral Pathology and Oral Radiology, 125(6), 582–590. 10.1016/j.oooo.2018.03.011

17. Weinstein, J. N., Collisson, E. A., Mills, G. B., Shaw, K. R. M., Ozenberger, B. A., Ellrott, K., Shmulevich, I., Sander, C., & Stuart, J. M. (2013). The Cancer Genome Atlas Pan-Cancer analysis project. Nature Genetics, 45(10), 1113–1120. 10.1038/ng.2764

18. Yu, H., Kim, P. M., Sprecher, E., Trifonov, V., & Gerstein, M. (2007). The Importance of Bottlenecks in Protein Networks: Correlation with Gene Essentiality and Expression Dynamics. PLoS Computational Biology, 3(4), e59. 10.1371/journal.pcbi.0030059

